# First Whole-Genome Chikungunya Virus Sequence Detected in Mosquitoes during the 2025 Foshan Outbreak: Evidence of Field Vector Infection and Transmission Potential in China

**DOI:** 10.1101/2025.08.25.672256

**Authors:** Xinyu Zhou, Xiaoxue Xie, Wenhao Wang, Heting Gao, Kai Wang, Xiaohui Liu, Xiaoli Chen, Yuting Jiang, Haotian Yu, Dan Xing, Teng Zhao, Chunxiao Li

## Abstract

**Background:** Since July 2025, an outbreak of mosquito-borne chikungunya fever occurred in Foshan City, Guangdong Province, China. This was the second outbreak in China following the one that occurred in Dongguan City, Guangdong Province, in 2010. Moreover, the intensity of this outbreak was significantly greater than that of the previous one. Updates to 23 August, more than 10,000 human cases had been reported. Here, we present the first full genome sequence of the chikungunya virus (CHIKV) derived from field-trapped mosquitoes during the outbreak.

**Methods:** Adult *Aedes albopictus* were BG-trap captured from residences and parklands in three hotspot towns with high density of confirmed human cases. Mosquitoes were morphologically identified and pooled by species, sex and environment types. RNA was extracted, screened by CHIKV RT-qPCR, then positive pools underwent sanger and whole-genome sequencing for complete sequences. Lineage and mutational profiles were inferred by maximum likelihood phylogenetic and comparison against human and mosquito genomes. The distribution of amino acid site mutations in different protein coding regions was also analyzed.

**Result:** Through 11 days of collection using 10 BG-traps, 2,803 mosquitoes were captured. 1569 (55.97%) female *Ae. albopictus* were divided into 77 pools and 9.09% (7/77) of the pools tested positive for CHIKV. The local *Ae. albopictus* minimum infection rate (MIR, per 1000 females) was 4.46, while the MIR for residences in Lecong Town was the highest at 9.17. The MIR for parklands was slight higher than for residences (4.60 vs. 4.30). All the 5 *Ae. albopictus*-derived complete CHIKV genome clustered within ECSA-Indian Ocean lineage genotype, closely related to human-derived genomes on 2025 Reunion Island. Amino-acid mutations E1-A226V/E2-L210Q were detected in the strains, which enhanced adaptability to *Ae. albopictus* and increased the transmission capacity. Novel mutation observed on E1 and E2 were totally consist to the patient-derived CHIKV in 2025 Reunion Island.

**Conclusions:** It was the first mosquito-derived CHIKV whole-genome during the 2025 Foshan outbreak, filling a critical gap between human case and entomological surveillance. *Ae. albopictus* was confirmed as the primary vector during the outbreak. The current outbreak CHIKV strain with particular amino-acid mutations had adapted to *Ae. albopictus* transmission. Compared to previous Chikungunya outbreaks over the past decade, the Foshan outbreak occurred earlier (early July), in a larger urban area (with a population of over 9.5 million), and with abundant breeding sites for the vector mosquito *Ae. albopictus*. However, the outbreak was quickly brought under control, with daily case numbers consistently decreasing, which is closely linked to the strong vector control measures implemented by the Chinese government in the early stages of the outbreak. Moreover, this event once again underscores the necessity of early monitoring of vector mosquitoes and the importance of implementing highly effective vector intervention measures as soon as possible after an outbreak occurs.

## 1. Introduction

The chikungunya virus (CHIKV) is a mosquito-borne single-stranded RNA virus (belonging to the *Togaviridae* family, *Alphavirus* genus) that causes an acute febrile illness accompanied by severe and debilitating arthralgia [1]. The primary vectors for CHIKV are *Aedes aegypti* and *Ae. albopictus*[2]. Given the widespread distribution of *Ae.* mosquitoes in tropical and subtropical regions, CHIKV has demonstrated significant global cross-regional transmission potential[3]. Currently, CHIKV is classified into four major genotypes: West African (WA), East/Central/South African (ECSA), Asian, and the Indian Ocean Lineage (IOL, a branch of the ECSA genotype). From the first detection from Tanzania in 1952, CHIKV was sporadically confined to Asia and Africa. However, since the beginning of the 21st century, it had re-emerged in over 100 countries across Asia, Africa and America. For example, the introduction of IOL had led to explosive epidemics in India and Southeast Asia, posing a serious threat to public health[4–6].

In 2010, Dongguan City in Guangdong witnessed China’s first large-scale local CHIKV outbreak, where abundant *Ae. albopictus* populations. Dongguan CHIKV strain belonged to the ECSA-IOL genotype with the E1-A226V mutation, which significantly enhances its adaptability to the local primary vector *Ae. albopictus* [7, 8]. Subsequently, imported cases and sporadic local transmission were reported in Yunnan, Zhejiang, and other regions[9, 10]. Nowadays, the emerging outbreak in Foshan City, Guangdong Province, which began in July 2025, had infected above 10 thousand people by August 23[11]. The subtropical climate and rapid urbanization in China, combined with high *Ae.* mosquito activity and population density, amplified the risk for CHIKV outbreaks in Pearl River Delta region (Guangdong province).

Viral genomes was instrumental in the identification of the epidemics origin and the reconstruction of transmission chains[12]. However, the majority of CHIKV viral sequences from the Foshan 2025 outbreak were patient-derived, with no field mosquito-derived CHIKV genome has been reported, which hindered comprehensive understanding of the transmission ecology and the establishment of phylogenetic linkage between human cases and local vector. There were several documented mosquito-derived CHIKV genomes from epidemics in Asia, European and Latin America countries[13–16]. Mosquito-derived CHIKV genomes not only validate vector infection but also reveal potential adaptive mutations. The E1-A226V mutation, first identified in the IOL strains of the 2005–2006 Indian Ocean outbreak, has been shown to dramatically increase CHIKV infectivity and dissemination in *Ae. albopictus*. In 2009, a second-step mutation (E2-L210Q) emerged in India, further enhancing midgut infection in *Ae. albopictus* without affecting fitness in *Ae. aegypti*. These sequential adaptations havd enabled CHIKV to exploit *Ae. albopictus* as a more efficient urban vector across a wider geographic and climatic range, where *Ae. albopictus* predominates, such as in urban China[17, 18].

This study was conducted during the ongoing CHIKV outbreak in August 2025, Foshan, China, where the CHIKV positivity rate was detected in *Ae.* mosquitoes and first mosquito-derived CHIKV strains whole genome was obtained. By assessing mosquito infection rates, genotypes, and key mutation profiles, this study provided critical evidence for CHIKV vector attribution and an evaluation of the mosquito control efficacy during the outbreak, enabling timely responses and guidance for mosquito-borne disease control measures.

## 2. Materials and Methods

### 2.1. Sampling Sites

This study was conducted in the field area of Beijiao, Chencun and Lecong Towns in Shunde District, Foshan City, Guangdong Province, China, which were a hotspot region (over 90% of patient cases were reported at the initial phase) for the CHIKV pandemic (Figure 1). The area is located approximately 30–40 kilometers from Guangzhou City (international city with 19 million population) and has a subtropical humid climate with an annual average temperature of 22.2°C (range 10°C to 33°C). The region receives approximately 1,677.3 millimeters of precipitation annually, with June having the highest precipitation at 273.7 millimeters and an annual average relative humidity of 79%, which is highly suitable for the survival of the primary vector *Ae.* mosquitoes.

**Figure 1.**
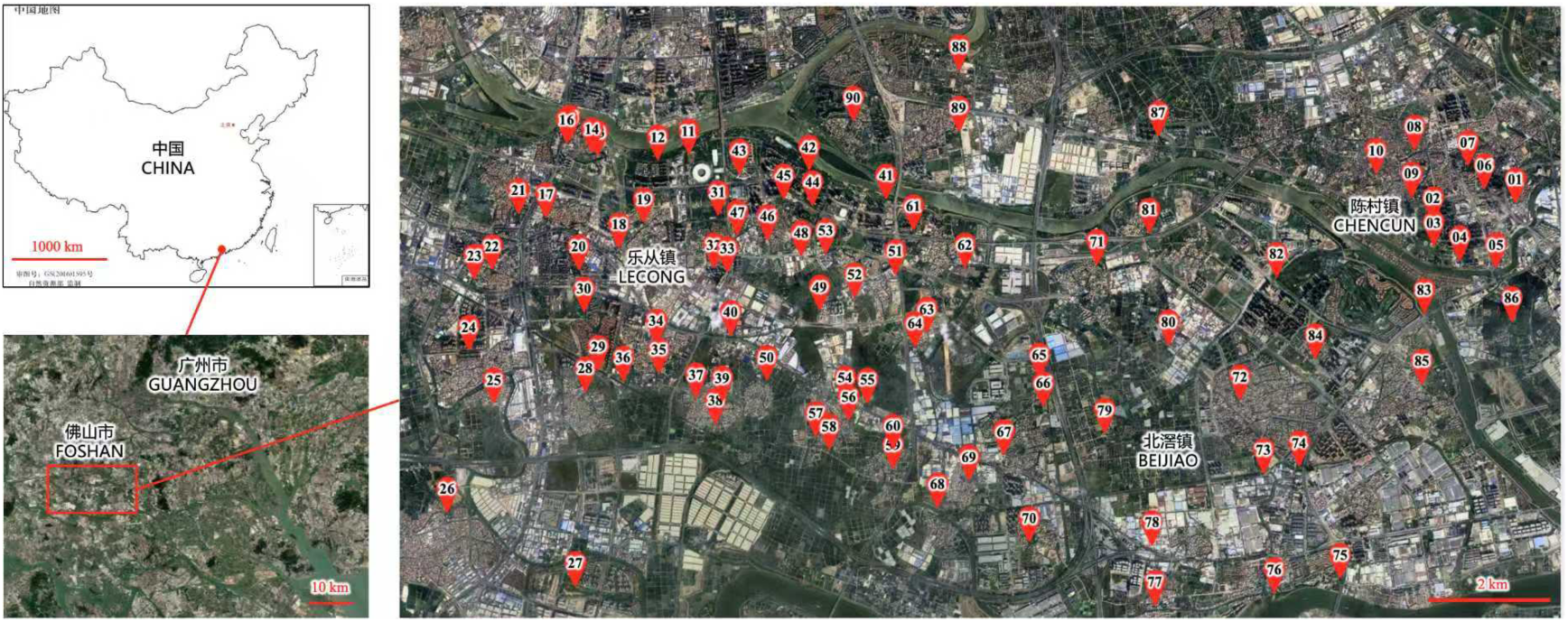
satellite map for sampling sites. Red points represented 90 sites in Beijiao, Chencun and Lecong towns, Foshan city, Guangdong Province, China

### 2.2. Mosquito Collection and processing

The mosquitoes were collected outdoors with BG-Sentinel traps from 9 am to 8 pm during July 31st and Aug 10th. Adult mosquitoes were frozen at − 20 °C for 30 min and placed on ice for morphological identification, then immediately transferred in carbon dioxide ice.

We restricted analyses to female *Ae.* mosquitoes, as only females blood-feed and contribute to arbovirus transmission. the *Ae. albopictus* mosquitoes were quantified and grouped in pools, according to species, sex, location. Each pool comprised approximately 25 individuals. Each pool was homogenized by motor driven tissue grinder with 1 ml of Media Dulbecco’s Modified Eagle Medium (DMEM) (Gibco) supplemented with 2% fetal bovine serum (FBS) (Gibco). Following homogenization, the samples were centrifuged at 8000 × g for 10 min at 4°C. The supernatant was subsequently collected and stored at -80°C until further processing.

### 2.3. Viral RNA Extraction and CHIKV molecular detection with RT-qPCR

The prepared homogenate was clarified by centrifugation and the supernatant was used for viral RNA isolation. RNA was extracted from pools of clarified mosquito homogenates using viral RNA isolation kit (QIAGEN, Germany, catalog #52906), according to manufacturer’s instructions. The RNA was eluted from the QIAspin columns in a final volume of 80μl of ddH20 and was kept at −80 ℃ until processing.

The RT-qPCR assay was performed using the CHIKV Detection Kit (SLin, China, catalog # A5-10B), which contains specific primers and a probe targeting the CHIKV E1 gene. Each 25 µL reaction contained 20 µL of reaction mix and 5 µL of extracted RNA. Amplification was carried out on an Applied Biosystems® QuantStudio™ 7 Flex Real-Time PCR System (ThermoFisher Scientific, Waltham, MA, USA) under the following conditions: reverse transcription at 50 °C for 2 min, initial denaturation at 95 °C for 1 min, followed by 40 cycles of denaturation at 95 °C for 2s and annealing/extension at 55 °C for 19s. A plasmid containing a fragment of the CHIKV E1 gene was used as a positive control, and nuclease-free water was included as a negative control. All samples were run in duplicate, and those with an average cycle threshold (Ct) value below 37 were considered positive.

### 2.4 Sanger sequencing of mosquito-derived CHIKV genomes

Full-length CHIKV genomes were amplified using overlapping RT-PCR with primer sets designed based on S27 strain reference sequences from GenBank (accession GCA_000854045.1). Full viral genome recovery was achieved via synthesis of complementary DNA (cDNA) directly from single-stranded RNA (ssRNA). Briefly, cDNA was synthesized using PrimeScript™ RT Master Mix (TAKARA, #RR036A) The CHIKV genome was sequenced through overlapping PCR amplicons spanning 12 genomic segments. All primers used for amplification (Table 1) were commercially synthesized by Sangon Biotech (Shanghai, China), with subsequent PCR product sequencing performed by the same vendor. Raw chromatograms were assembled into contiguous sequences using SeqMan Pro (DNASTAR Lasergene v7.1) and manually curated to generate the complete genome sequences.

**Table 1.**
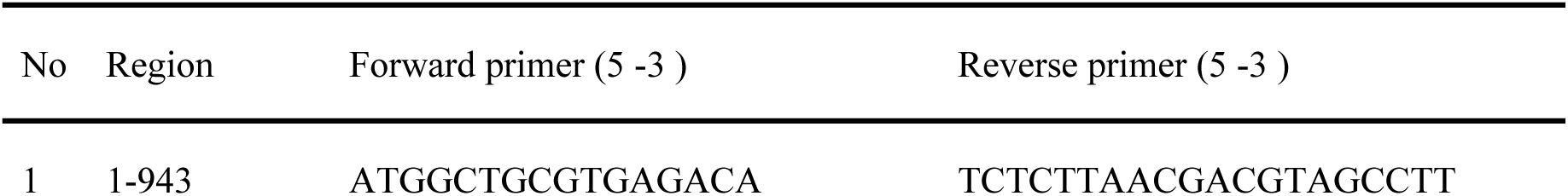
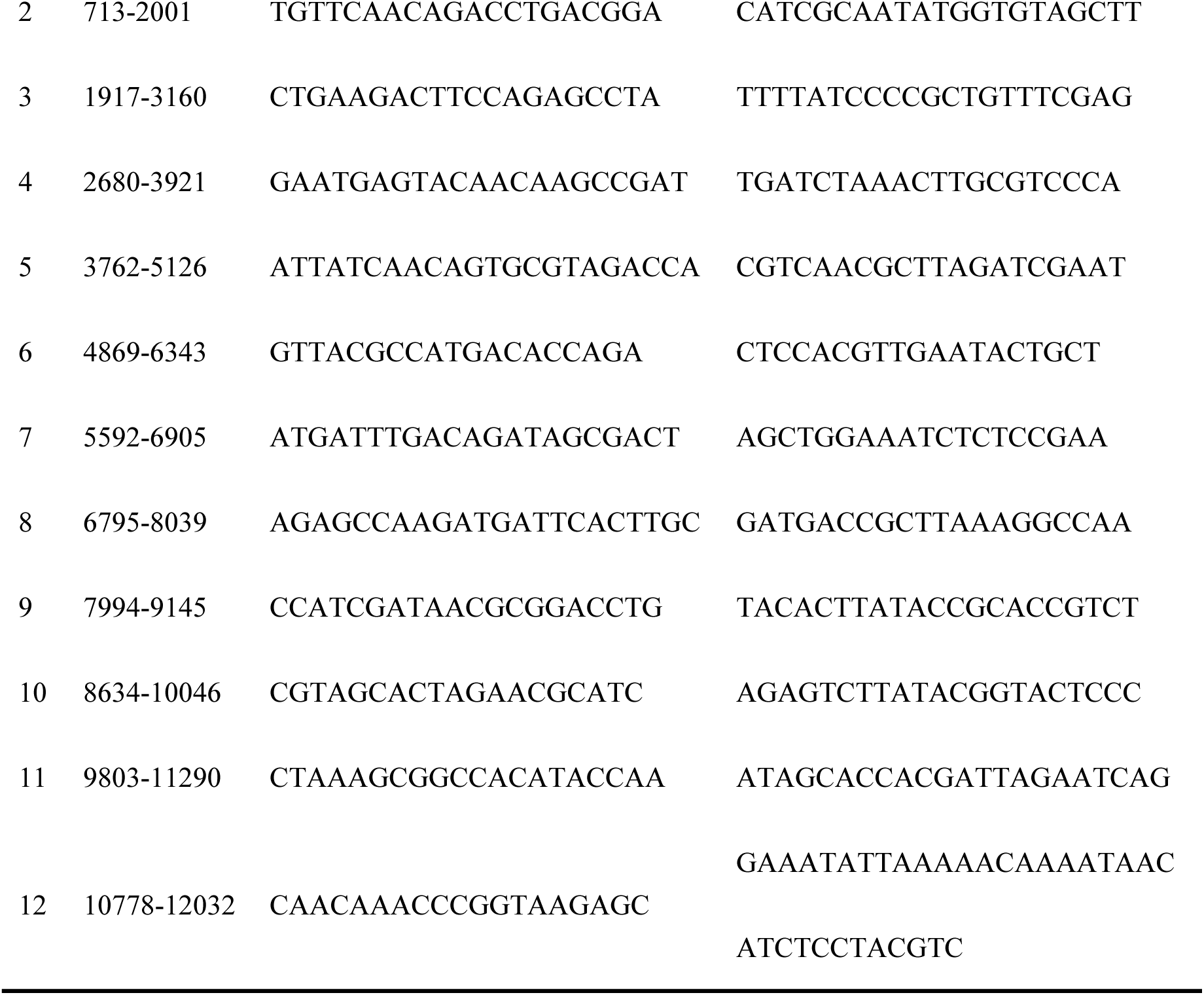
Primers for sanger sequencing of mosquito-derived CHIKV genomes.

### 2.5. Next-generation sequencing of mosquito-derived CHIKV genome

Due to the low viral RNA concentration, some PCR-positive pools were subjected to high-throughput sequencing. cDNA libraries were prepared using the NEBNext Ultra RNA Library Prep Kit for Illumina. The libraries were quality checked and sequenced on an Illumina Nova Seq 6000 platform with 150-bp paired-end reads.

The analysis pipeline for high-throughput sequencing data was as follows: First, Trimmomatic v0.39[19] and Fastp v0.23.4[20] were used to remove sequencing adapters and low-quality sequences, completing the sequencing data quality control. Next, Bowtie2 v2.5.4[21] was employed in conjunction with the *Ae. albopictus* reference genome (Accession Number: GCF035046485.1)[22] to perform host sequence depletion on the quality-controlled sequencing results. Subsequently, the host-depleted sequences were aligned against the Chikungunya virus genome sequence (Accession Number: NC_004162.2)[23] using ncbi-blast+ v2.16.0[24], and Chikungunya virus-specific sequences were isolated from the alignment output. These isolated viral sequences were then assembled with MEGAHIT v1.2.9[25]. For the assembled contigs, BWA v0.7.17-r1188[26] was used to evaluate the sequencing coverage depth, and ncbi-blast+ v2.16.0 for sequence homology alignment. Finally, the final high-quality assembly results were screened and obtained through these validation steps.

### 2.6 Phylogenetic Analysis

Besides the 5 mosquito-derived CHIKV strains detected in Foshan, an additional 125 complete genomes of Chikungunya virus were collected and curated, with isolation sources including mosquitoes and humans. It should be noted that, as the sudden outbreak, no sequences from local patients in Foshan 2025 had been found in the database. Following codon-based multiple sequence alignment using MUSCLE v3.8.1551[27], the Bayesian Evolutionary Analysis Utility (BEAUti) v10.5.0[28] was used for evolutionary model selection and XML file construction.

Bayesian Evolutionary Analysis Sampling Trees (BEAST) v10.5.0[29] was employed to estimate evolutionary rates, divergence times, population sizes, and tree topologies. Tracer v1.7.2[30] was utilized to assess the convergence of phylogenetic tree topology parameters.

TreeAnnotator v10.5.0[31] was used to summarize the posterior estimates and highest posterior density (HPD) limits of node heights, as well as evolutionary rates for analyses employing a relaxed molecular clock model. Finally, FigTree v1.4.4 was applied to visualize and refine the phylogenetic results for presentation purposes.

### 2.7 Amino acid mutation analysis

For sequence variant detection, the complete genomes of each Foshan mosquito-derived CHIKV strain were aligned to the typical genotypes from human and mosquito reference strains using ClustalW[32], and the bases of each Foshan mosquito-derived CHIKV strain virus genome that did not align were extracted as single-nucleotide variants (SNVs) using custom-written Python scripts. These SNVs were annotated by ANNOVAR software, SNVs in the coding region were divided into synonymous SNVs and nonsynonymous SNVs.

### 2.8 Comparative genomic analysis

Predict the sequences and genomic positions of structural and non-structural proteins of Chikungunya virus using blastn, and plot the genome structure using R package circlize v0.4.16[33].

### 2.9 Minimum infection rate

For infection rate estimation, we restricted analyses to female *Ae.* mosquitoes, as only females blood-feed and contribute to arbovirus transmission. Pools were formed with around 25 individuals, and infection rates were expressed as minimum infection rate (MIR, per 1,000 females). The MIR was determined as the ratiobetween the number of virus positive pools of mosquitoes detectedand the total number of mosquitoes tested, multiplied by 1000. This approach has been widely adopted in arbovirus entomological surveillance to avoid dilution effects from non-vector males, which may bias estimates downward.

## 3. Results

### 3.1. Mosquito Collection and Identification

Sampling sites were taken from three hotspot towns (Beijiao, Chencun and Lecong) in Foshan city, Guangdong Province, China, including 56 parklands and 34 residential areas. A total of 2,803 mosquitoes (1031 in Beijiao, 500 in Chencun and 1272 in Letang) were trapped in the study. The majority were *Ae. albopictus* (2,627, or 93.72%), with the others being Culex spp., with no *Ae. Aegypti.* There were 1087 female *Ae. Albopictus* trapped from parklands and 465 from residences. (Supplementary 1).

### 3.2 CHIKV infection rate in mosquito

We put the focus on female *Ae.* mosquitoes, as only females blood-feed and contribute to arbovirus transmission. There were 1569 (59.73%) female *Ae. Albopictus* from 90 sites in 3 towns, which were divided into 77 pools by species, gender and Environmental type. There were 9.09% (7/77) of the pools tested positive for CHIKV (Table 2).

**Table 2.**
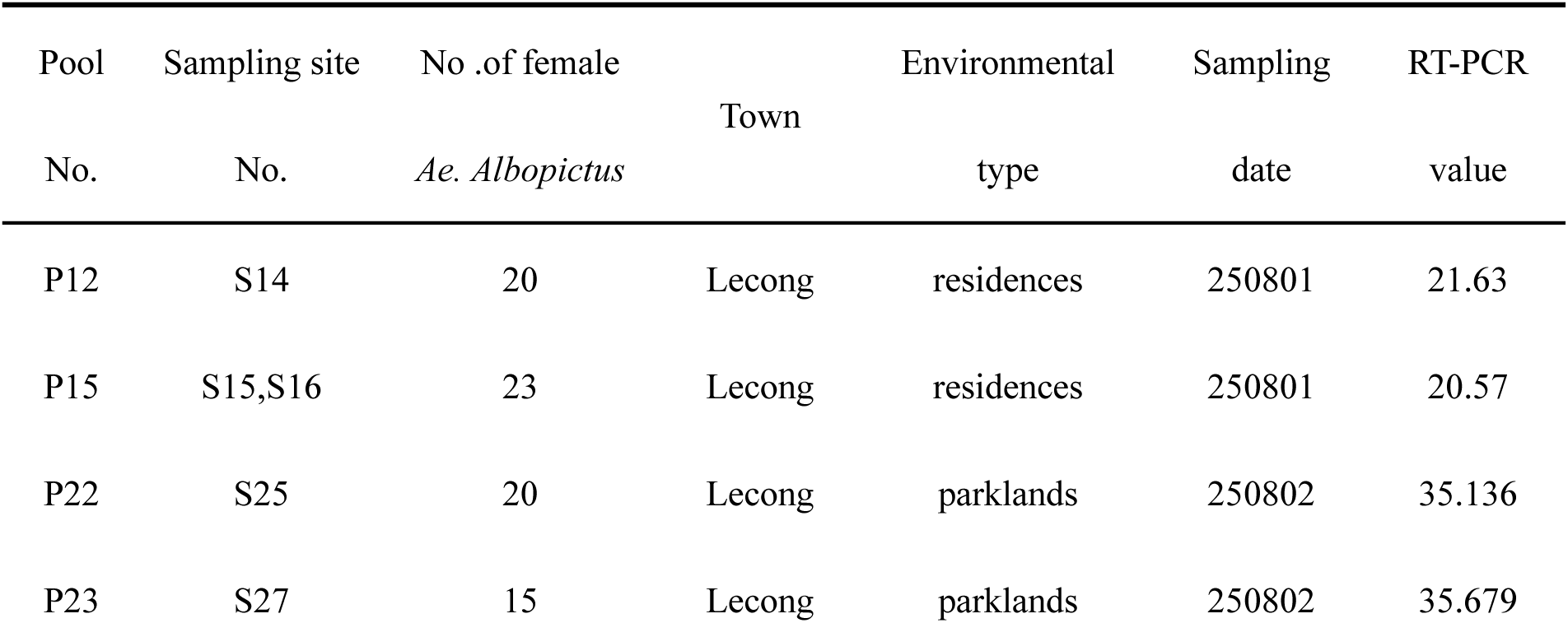
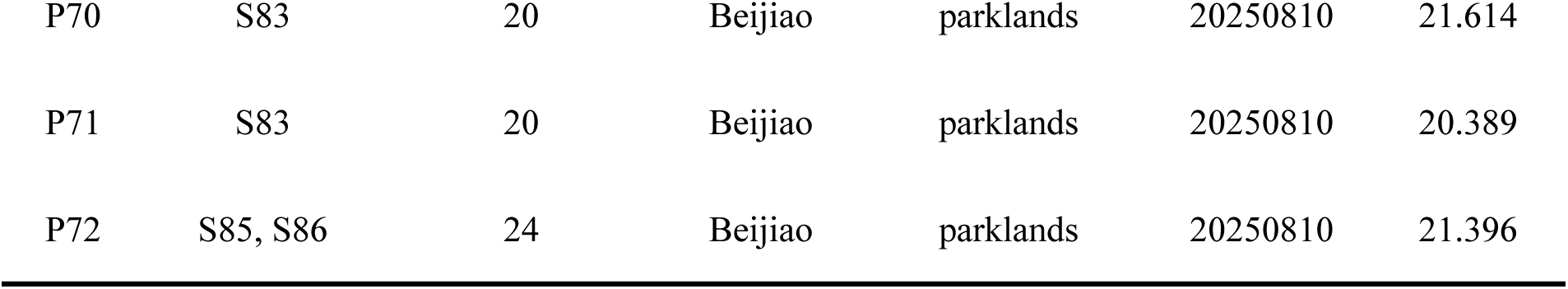
CHIKV Positive pool information for female *Ae. albopictus* mosquitoes.

| Pool No. | Sampling site No. | No .of female *Ae. Albopictus* | Town | Environmental type | Sampling date | RT-PCR value |
| --- | --- | --- | --- | --- | --- | --- |
| P12 | S14 | 20 | Lecong | residences | 250801 | 21.63 |
| P15 | S15,S16 | 23 | Lecong | residences | 250801 | 20.57 |
| P22 | S25 | 20 | Lecong | parklands | 250802 | 35.136 |
| P23 | S27 | 15 | Lecong | parklands | 250802 | 35.679 |
| P70 | S83 | 20 | Beijiao | parklands | 20250810 | 21.614 |
| P71 | S83 | 20 | Beijiao | parklands | 20250810 | 20.389 |
| P72 | S85, S86 | 24 | Beijiao | parklands | 20250810 | 21.396 |

The total minimum infection rate (MIR, per 1000 females) was 4.46, while the MIR for residences in Lecong Town was the highest at 9.17. The MIR for parklands was slight higher than for residences (4.60 vs. 4.30, Table 3).

**Table 3.** Minimum infection rate (MIR) of CHIKV in mosquito from different towns and regional types.

| Town | Environmental<br>type | Total of female<br>*Ae. Albopictus* | Total of<br>the pools | No. of pools<br>positive by RT-PCR (%) | MIR per 1000<br>female |
| --- | --- | --- | --- | --- | --- |
| Beijiao | parklands | 437 | c21 | 3 (14.28) | 6.86 |
|  | residences | 162 | 8 | 0 | 0 |
| Chencun | parklands | 217 | 11 | 0 | 0 |
|  | residences | 85 | 4 | 0 | 0 |
| Lecong | parklands | 450 | 22 | 2 (9.09) | 4.44 |
|  | residences | 218 | 11 | 2 (18.18) | **9.17** |
| Total | parklands | 1087 | 54 | 5 (9.26) | 4.60 |
|  | residences | 465 | 23 | 2 (8.70) | 4.30 |
|  | Total | 1569 | 77 | 7 (9.09) | **4.46** |

### 3.3. Phylogenetic Analysis

There were 5 whole genome sequence finally acquired, from the PCR positive pools excluded 2 low viral load pools (CT value beyond 35). The initial Maximum clade credibility tree was constructed using the dataset containing 135 sequences from the four distinct genotypes and the Foshan mosquito-derived sequences in the study. Phylogeny based on the complete genome analysis characterized the 5 Foshan mosquito-derived CHIKV strains (12/15/70/71/72_Guangdong_foshan_Mosquito_2025), as belonging to the ECSA-IOL genotype, with a high similarity to the Réunion Island human case in 2025 (99.93%) and local human case in 2018 (97.01%∼97.04%). It comprised a distinct sub-branch with other mosquito derived samples previously detected in Guangdong, Zhejiang and Yunnan (Figure 2). This showed a potential genetic origin of the virus strains in this outbreak.

**Figure 2.**
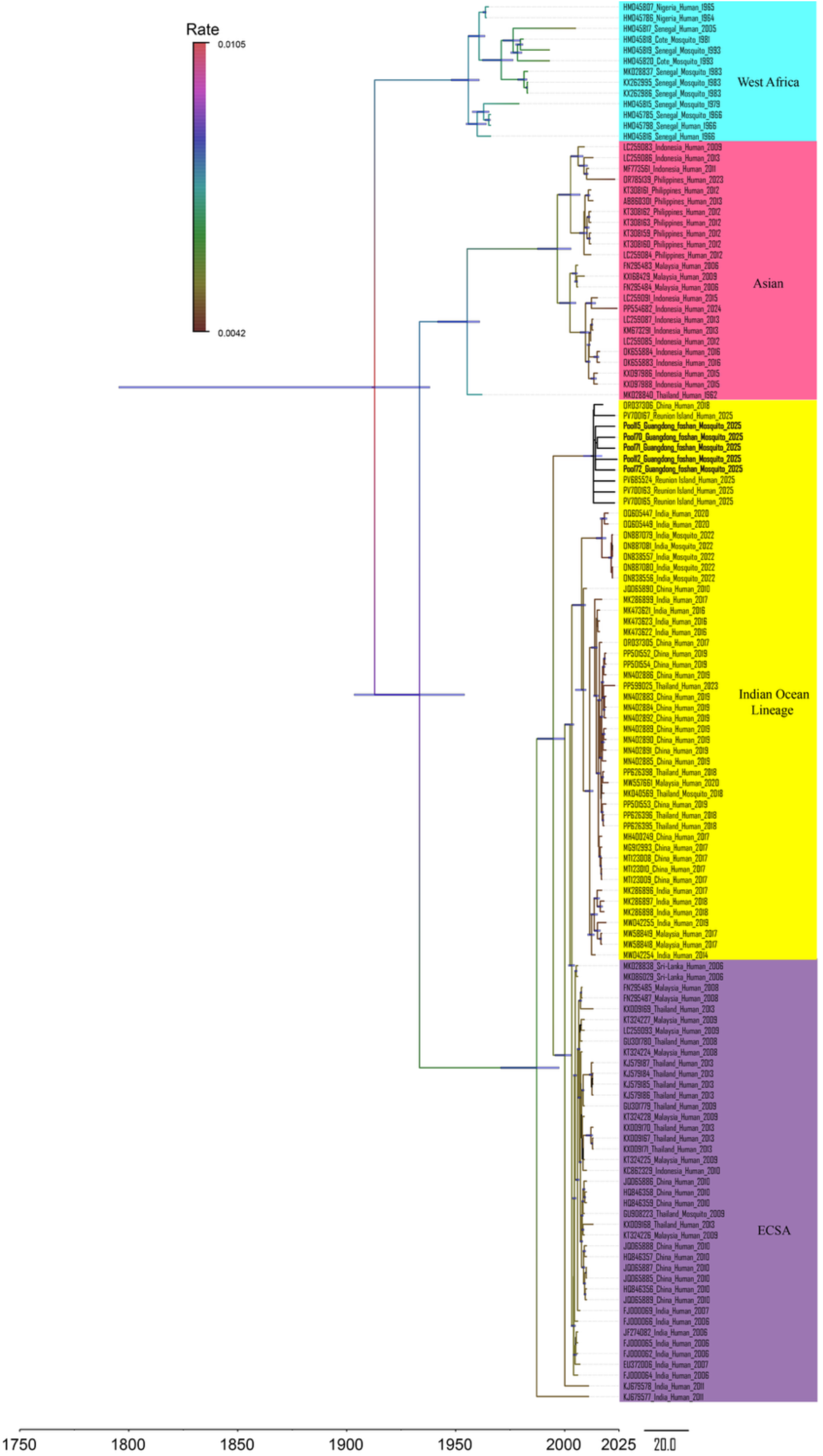
Maximum clade credibility (MCC) tree of 130 CHIKV strains. The four major lineages are highlighted with different branch colors, where the color of each branch line represents the evolutionary rate of the viruses in that branch. The estimated 95% HPD values for most pecent common ancestors are labeled beside the node and are also indicated by the thick blue horizontal node bars. The numbers adjacent to nodes indicate Bayesian posterior probability values. Strains are labeled as follows: strain_name_location_host_date (year) of collection. The 5 Foshan mosquito-derived CHIKV strains in this study were in bold.

Phylogenetic results indicated that all currently circulating CHIKV strains share a common ancestor that existed within the past 300 years, with the 95% highest posterior density (HPD) interval for their most recent common ancestor (MRCA) estimated to be 87 – 230 years ago. For CHIKV strains endemic to the Asian region, the 95% HPD interval for their MRCA ranges from 71 to 122 years ago. The divergence between the Asian and ECSA genotypes occurred within the past 100 years. Furthermore, phylogenetic results revealed a distinct spatiotemporal pattern in the Southeast Asian lineage of the Asian genotype: it spread from Thailand to Indonesia, subsequently to the Philippines, and most recently to Malaysia. The most recent common ancestor (MRCA) of the Indian Ocean lineage can be traced back to approximately 2002 (95% HPD: 2001– 2003).

### 3.4 Amino acid mutation analysis

The mutation analysis, compared to human- and mosquito-derived strains, was shown in Table 4 for four genotypes of CHIKV structural proteins. All consensus genomes assigned to the five Foshan 2025 mosquito-derived CHIKV strains (Pools 12, 15, 70, 71 and 72) displayed an identical E1/E2 amino-acid signature, indicating a single predominant circulating variant in local vectors.

**Table 4.**
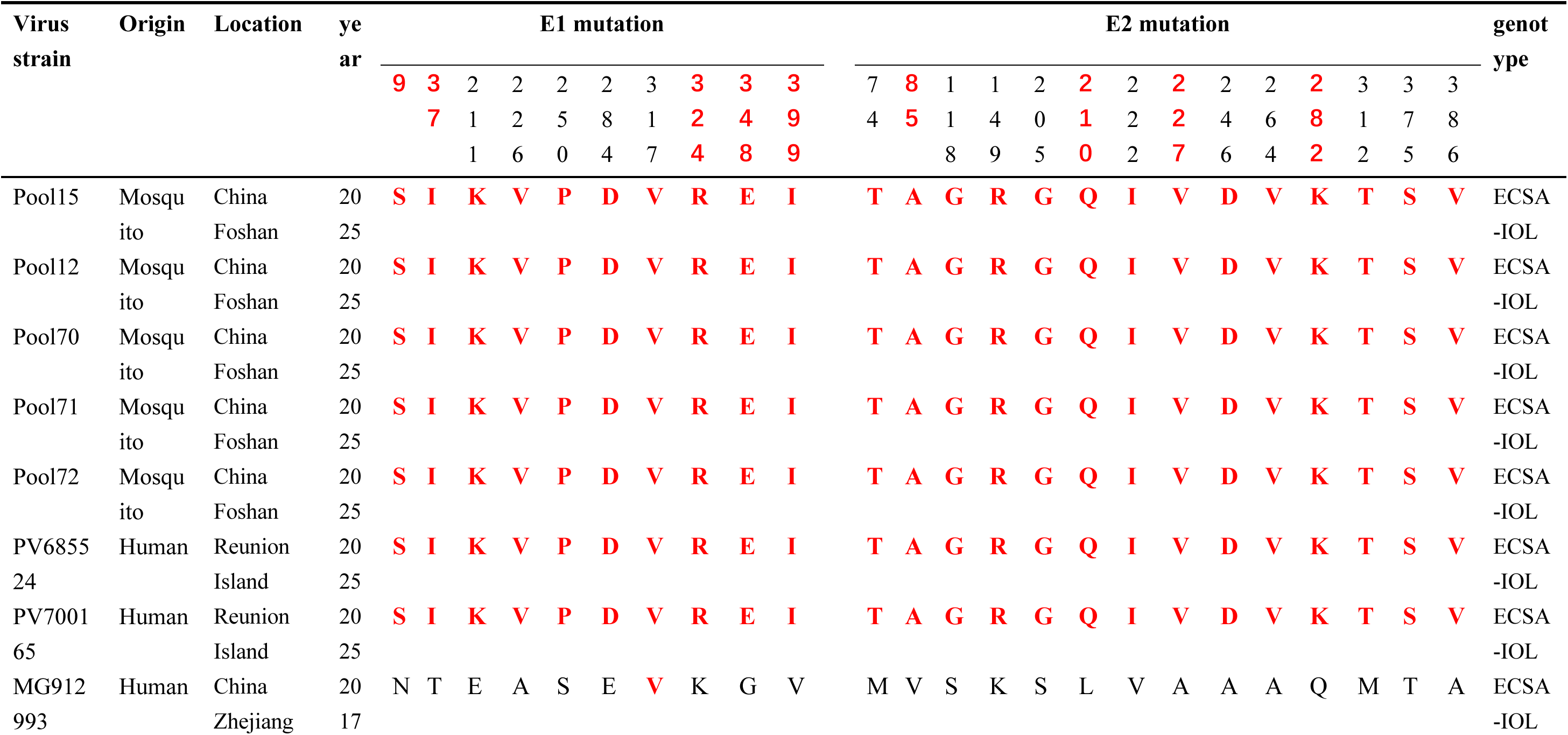
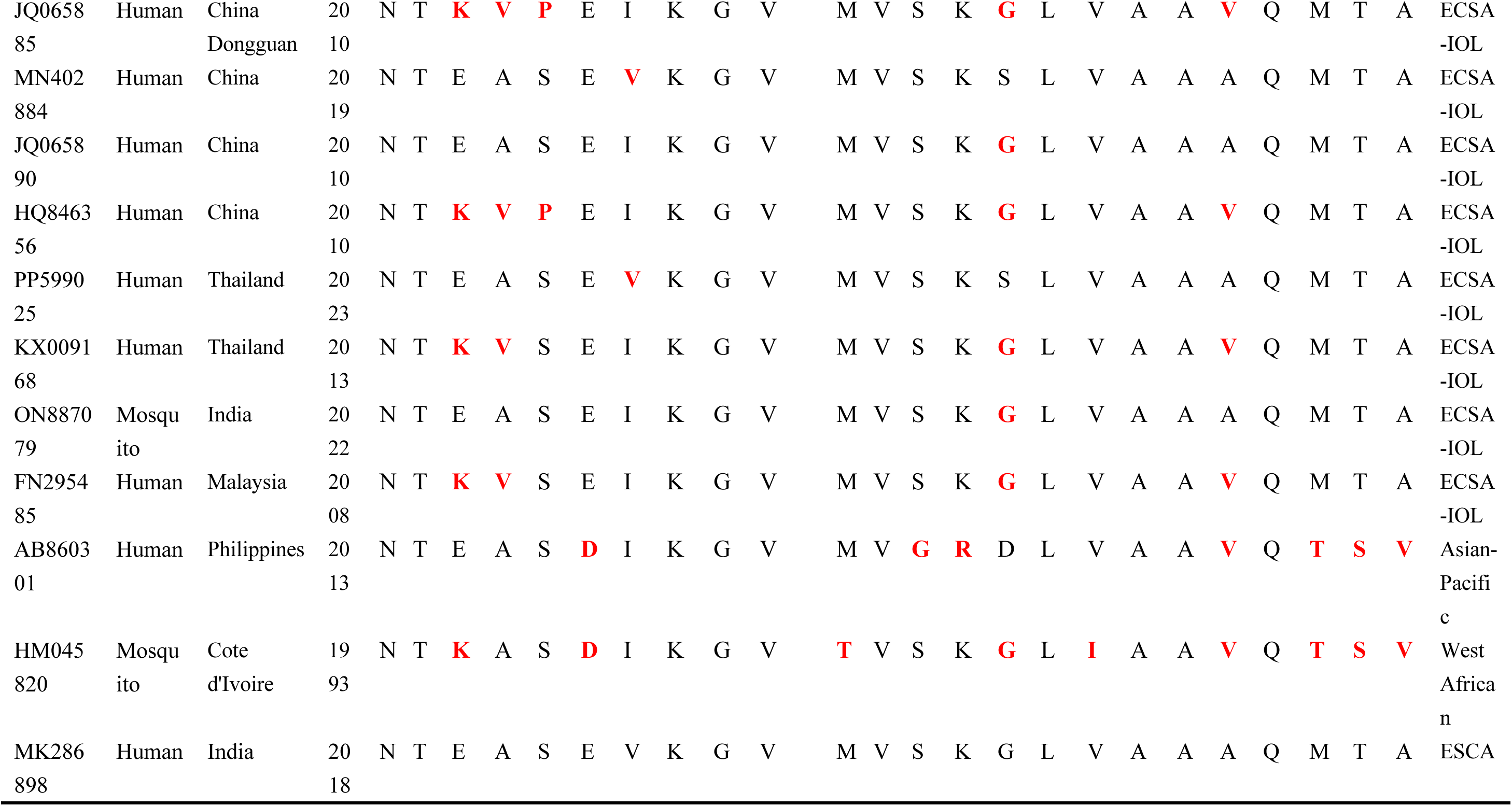
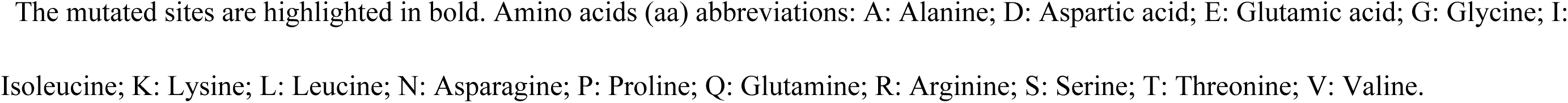
Mutations in Foshan mosquito-derived CHIKV strains, compared with typical sequences from human and mosquito in the four genotypes.

Notably, this constellation included the E1-A226V and E2-L210Q mutations, which had been identified as enhancing *Ae. albopictus* adaptation and increasing infectivity. The novel E1-N9S/T37I/K324R/G348E/V399I mutation together with the E2-V85A/A227V/Q282K mutation were observed, which were identical to those observed in two contemporaneous Réunion Island human isolates (PV685524 and PV700165) in 2025, supporting a shared variant profile across regions and facilitating tracing.

Contrasting with earlier Chinese reference strains (e.g. Dongguan 2010 and Zhejiang 2017), the Foshan variant consistently replaced ancestral residues at multiple E1 sites (N9S, T37I, K324R, G348E and V399I) and E2 sites (V85A, A227V and Q282K). These changes made the 2025 Foshan mosquito-derived CHIKV different from former domestic lineages within ECSA–IOL.

### 3.5 Comparative genomic analysis

BLAST analysis and nonsynonymous SNV were used to illustrate five Foshan mosquito-derived CHIKV genomes. Furthermore, the complete sequencing of the 5 Foshan isolates and other 20 sequnences from patients and mosquitos were collected, allowed for a better understanding of their genetic relationships. The different regions of the sequences exhibited > 90.4% nucleotide similarity to the corresponding regions of the prototypical isolate (Figure 3).

**Figure 3.**
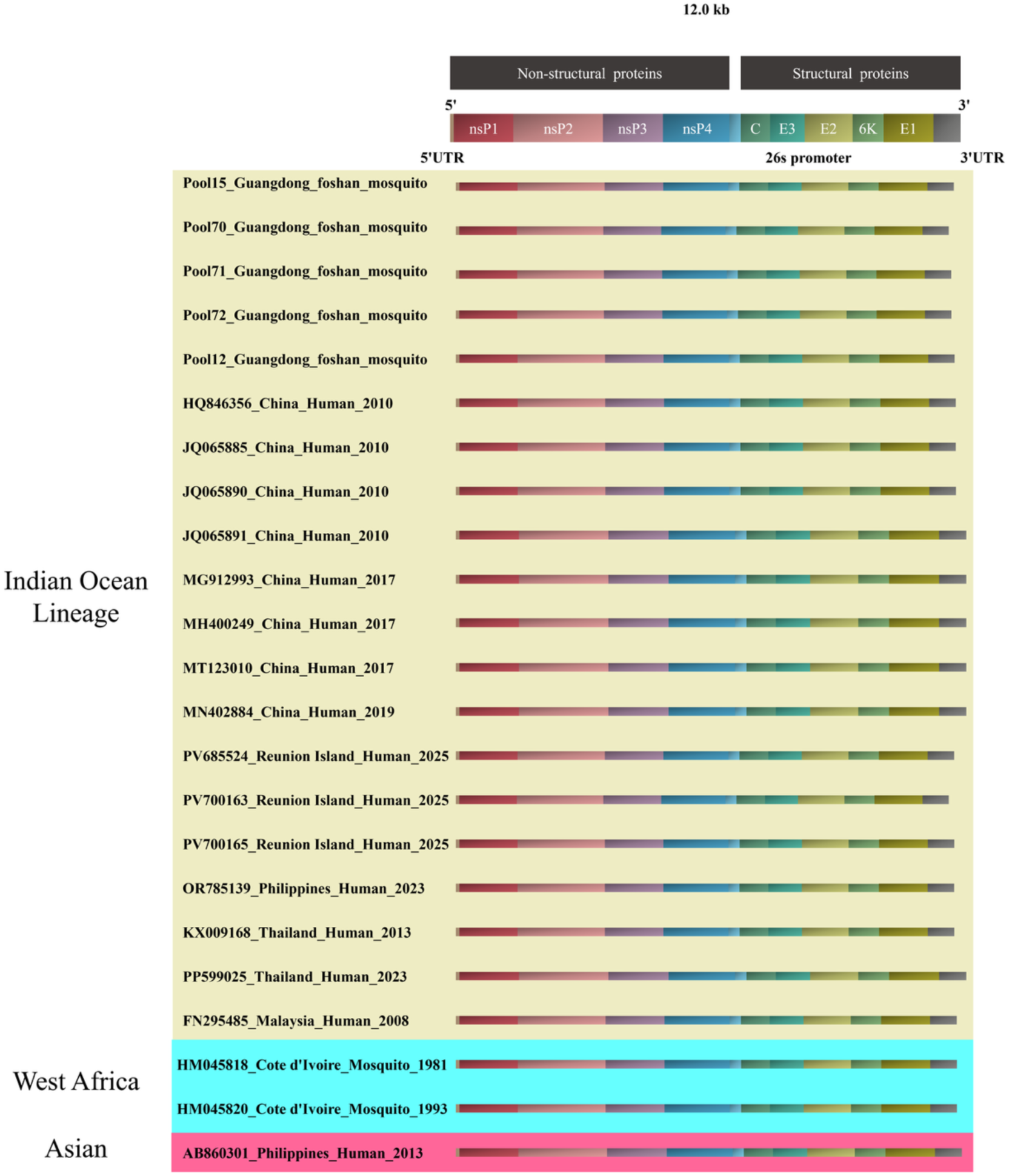
Comparative genomics analysis of 25 CHIKV strains. It showed the positions of gene-coding regions across different CHIKV lineages. Here, nsP stands for non-structural proteins, E for Envelope proteins, C for Capsid protein, and UTR for Untranslated Regions. Different background colors are used to distinguish between different CHIKV genotypes.

## 4. Discussion

This study reported the first mosquito-derived complete CHIKV genome from *Ae. albopictus* during the ongoing 2025 Foshan China outbreak, providing direct evidence of the vector mosquito *Ae. albopictus*‘s role in the CHIKV transmission. The observed mosquito infection rate (MIR = 4.46), together with adaptive mutations such as E1-A226V and E2-L210Q, provided a timely and effective support for public health response.

Globally, CHIKV detection rate in *Ae.* mosquitoes consistently increased during epidemic periods compared with inter-epidemic phases. For instance, a large-scale epidemic in India from 2006 to 2010 reported the MIR of 2-15 per 1,000 *Ae. aegypti* and *Ae. albopictus*, while while routine surveillance typically detected <1 per 1,000[34, 35]. Studies from Thailand, Indonesia, Singapore and other Asia countries, similarly demonstrated significant increases in *Ae. albopictus* positivity rates during outbreaks, reaching >8% in some settings, compared to sporadic detections in non-outbreak years[36–38] . The situation for *Ae. aegypti* was quite similar. Latin American surveys, such as in Colombia (2020– 2021), revealed very low positivity in *Ae. aegypti* during non-outbreak periods (0.2%), whereas epidemic periods in Brazil (2017–2020) reached 2–8 per 1,000[39, 40]. Higher infection levels were reported in Africa, like Kenya exceeded 10/1000[41]. Collectively, these findings confirmed that vector positivity rates rise sharply during outbreaks, and the Foshan outbrek MIR falls within the range observed elsewhere, suggesting comparable transmission dynamics and the efficacy of existing control measures

Phylogenetic analysis showed that the five Foshan mosquito-derived CHIKV genomes clustered with the Réunion Island human case in 2025, belonging to the same ESCA-IOL genotype as Foshan Local human-derived CHIKV did[11]. Besides, the Foshan mosquito-derived CHIKV amino-acid mutation on E1/E2 regions were identical to those observed in Réunion Island human isolates in 2025, supporting a potential genetic origin. Importantly, mutations such as E1-A226V and E2-L210Q, both detected in this study, were documented markers of enhanced vector adaptation. E1-A226V was known to increase infectivity in *Ae. albopictu*s approximately 100-fold by facilitating midgut invasion and shortening the extrinsic incubation period[42]. Similarly, the E2-L210Q allele had been associated with enhanced viral replication efficiency within the vector[43]. These mutations, first highlighted during the 2005– 2006 Indian Ocean epidemic, have since emerged independently in multiple regions, like Thailand and Malaysia [38, 44], reflecting selective pressure for *Ae. albopictus* adaptation. Conversely, outbreaks in the Americas had largely involved Asian genotype viruses lacking E1-A226V, demonstrating regional variation in adaptive signatures[45]. The Foshan mosquito-derived CHIKV strains in this study carried the E1-A226V and E2-L210Q mutations, which were of significant importance for explaining the CHIKV adaptive transmission in local primary vector *Ae. albopictus*. Other E2 substitutions such as R198Q, K233E, and K252Q have also been shown to provide incremental fitness benefits in *Ae. albopictus* within IOL strains[46, 47]. Other amino-acid changes observed in Foshan strains had not been previously implicated in vector adaptation so far.

Despite early implementation of large-scale vector control during the Foshan outbreak, CHIKV-positive mosquitoes were still detected, which was consistent with observations in Brazil, Thailand and other regions, where wild-caught mosquitoes remained virus-positive even under intensive interventions. This persistence likely reflected challenges such as complex mosquito habitats, insecticide resistance[48, 49]. Comparisons with the concurrent 2025 Reunion Island outbreak were instructive: although Reunion experienced lower seasonal mosquito abundance, nearly one-quarter of the population was infected, with >23,000 estimated cases per week[50, 51]. In Foshan, despite higher population density and favorable conditions for mosquito activity, timely interventions limited the cases to 10,000, underscoring the critical role of rapid, intensive vector management. Otherwise, the outbreak would have led to greater spread and loss.

Our findings emphasized that mosquito-based surveillance was indispensable for outbreak preparedness. Incorporating systematic human/mosquito monitoring and viral genome sequencing into public health responses provided an early warning system, informed on adaptive mutations, and enabled real-time evaluation of intervention efficacy. MIR estimates, when combined with genomic data, could guide adjustments to control intensity and complement case-based surveillance.

Compared to previous Chikungunya outbreaks over the past decade, the Foshan outbreak occurred earlier (early July), in a larger urban area (with a population of over 9.5 million), and with abundant breeding sites for the vector mosquito *Ae. albopictus*. However, the number of cases has so far just exceeded 10,000, and the number of daily new cases is under effective control, which is closely linked to the strong vector control measures implemented by the Chinese government in the early stages of the outbreak. Moreover, this event once again underscores the necessity of early monitoring of vector mosquitoes and the importance of implementing highly effective vector intervention measures as soon as possible after an outbreak occurs.

## Conclusions

This study provided the first complete CHIKV genome sequence obtained from mosquitoes during an ongoing endemic outbreak in China, bridging the existing gap between human and vector. Furthermore, the analysis of mosquito positive rates and amino-acid mutation spectra directly providing crucial support for the process of tracing and risk assessment. It emphasized the importance of corporating mosquito surveillance, in order to support a timely and effective public health response.

## Supplementary Information

Additional file 1 Mosquito collection information in 90 sampling sites, Foshan, China, 2025

## Ethics approval and consent to participate

Not applicable

## Consent for publication

All authors read and approved the final manuscript for publication.

## Availability of data and materials

Data and materials will be made available on request.

## Author contributions

Field mosquito collection: Xinyu Zhou, Xiaoxue Xie, Wenhao Wang; Sanger sequencing: Xiaoxue Xie, Xiaohui Liu, Xiaoli Chen; Whole-genome sequencing : Heting Gao; Developed the methodology: Dan Xing, Chunxiao Li; Collected the data: Kai Wang, Yuting Jiang, Haotian Yu; Analyzed the results: Heting Gao, Teng Zhao; Wrote the first draft: Teng Zhao; Generated the figures: Xinyu Zhou, Wenhao Wang; Conceptualization, resources and funding: Chunxiao Li. All authors read and approved the final manuscript.

## Acknowledgement

Not applicable

---

**Supplementary 1.** Mosquito collection information in 90 sampling sites, Foshan, China, 2025.

| Sampling site No. | longitude and latitude | Town | Environmental type | Sampling date | Mosquito collection |  |  |
| --- | --- | --- | --- | --- | --- | --- | --- |
|  |  |  |  |  | No. of mosquito | No. of *Ae. albopictus* | No. of Female *Ae. albopictus* |
| S1 | 22.955475,113.241928,-1.79 | Chencun | Parklands | 2025.07.31 | 30 | 26(0.87) | 17(0.65) |
| S2 | 22.953634,113.229721,1.17 | Chencun | Parklands | 2025.07.31 | 0 | 0(0) | 0(0) |
| S3 | 22.949833,113.230072,-2.82 | Chencun | Residences | 2025.07.31 | 36 | 34(0.94) | 15(0.42) |
| S4 | 22.947151,113.233315,-2.22 | Chencun | Residences | 2025.07.31 | 6 | 5(0.83) | 5(0.83) |
| S5 | 22.945978,113.238266,1.36 | Chencun | Parklands | 2025.07.31 | 51 | 51(1) | 44(0.86) |
| S6 | 22.957052,113.238411,-3.34 | Chencun | Parklands | 2025.07.31 | 85 | 84(0.99) | 19(0.22) |
| S7 | 22.961157,113.236382,3.09 | Chencun | Parklands | 2025.07.31 | 9 | 7(0.78) | 5(0.56) |
| S8 | 22.964167,113.227379,4.74 | Chencun | Parklands | 2025.07.31 | 37 | 30(0.81) | 5(0.14) |
| S9 | 22.957512,113.227165,-1.71 | Chencun | Parklands | 2025.07.31 | 21 | 10(0.48) | 5(0.24) |
| S10 | 22.959925,113.221344,-4.42 | Chencun | Residences | 2025.07.31 | 48 | 48(1) | 40(0.83) |
| S11 | 22.972282,113.112000,6.72 | Lecong | Parklands | 2025.08.01 | 160 | 160(1) | 59(0.37) |
| S12 | 22.972204,113.107788,2.21 | Lecong | Parklands | 2025.08.01 | 9 | 9(1) | 8(0.89) |
| S13 | 22.973234,113.097481,-0.84 | Lecong | Residences | 2025.08.01 | 22 | 16(0.73) | 5(0.23) |
| S14 | 22.973757,113.096970,-3.98 | Lecong | Residences | 2025.08.01 | 36 | 26(0.72) | 22(0.61) |
| S15 | 22.976496,113.093040,-0.01 | Lecong | Residences | 2025.08.01 | 74 | 74(1) | 54(0.73) |
| S16 | 22.975836,113.092888,-4.01 | Lecong | Residences | 2025.08.01 | 37 | 36(0.97) | 29(0.78) |
| S17 | 22.965343,113.089172,-2.75 | Lecong | Parklands | 2025.08.01 | 23 | 11(0.48) | 6(0.26) |
| S18 | 22.959852,113.100372,0.09 | Lecong | Parklands | 2025.08.01 | 45 | 42(0.93) | 32(0.71) |
| S19 | 22.963625,113.104378,-8.28 | Lecong | Parklands | 2025.08.01 | 13 | 10(0.77) | 6(0.46) |
| S20 | 22.956902,113.093819,1.90 | Lecong | Parklands | 2025.08.01 | 10 | 10(1) | 8(0.8) |
| S21 | 22.965977,113.085014,-1.06 | Lecong | Parklands | 2025.08.02 | 9 | 9(1) | 7(0.78) |
| S22 | 22.957771,113.080109,-1.66 | Lecong | Parklands | 2025.08.02 | 4 | 4(1) | 1(0.25) |
| S23 | 22.957047,113.077057,0.69 | Lecong | Parklands | 2025.08.02 | 14 | 14(1) | 9(0.64) |
| S24 | 22.946693,113.075287,-2.69 | Lecong | Residences | 2025.08.02 | 1 | 1(1) | 1(1) |
| S25 | 22.938643,113.079262,-1.06 | Lecong | Parklands | 2025.08.02 | 50 | 48(0.96) | 27(0.54) |
| S26 | 22.923597,113.070625,-1.71 | Lecong | Parklands | 2025.08.02 | 34 | 34(1) | 8(0.24) |
| S27 | 22.911486,113.090263,-3.43 | Lecong | Parklands | 2025.08.02 | 34 | 34(1) | 15(0.44) |
| S28 | 22.939260,113.093658,-4.03 | Lecong | Residences | 2025.08.02 | 1 | 1(1) | 1(1) |
| S29 | 22.941837,113.095718,- 10.78 | Lecong | Parklands | 2025.08.02 | 86 | 55(0.64) | 19(0.22) |
| S30 | 22.950142,113.094528,-11.59 | Lecong | Residences | 2025.08.02 | 22 | 22(1) | 12(0.55) |
| S31 | 22.963081,113.116425,0.54 | Lecong | Parklands | 2025.08.03 | 27 | 20(0.74) | 14(0.52) |
| S32 | 22.955696,113.115623,-5.01 | Lecong | Residences | 2025.08.03 | 6 | 6(1) | 5(0.83) |
| S33 | 22.955128,113.117287,-4.74 | Lecong | Parklands | 2025.08.03 | 31 | 31(1) | 19(0.61) |
| S34 | 22.945007,113.105316,-4.06 | Lecong | Residences | 2025.08.03 | 6 | 6(1) | 2(0.33) |
| S35 | 22.940918,113.105118,-4.17 | Lecong | Parklands | 2025.08.03 | 22 | 20(0.91) | 13(0.59) |
| S36 | 22.940413,113.099709,-2.79 | Lecong | Residences | 2025.08.03 | 45 | 45(1) | 24(0.53) |
| S37 | 22.936750,113.110977,-5.83 | Lecong | Residences | 2025.08.03 | 16 | 16(1) | 12(0.75) |
| S38 | 22.932940,113.113350,-5.53 | Lecong | Parklands | 2025.08.03 | 14 | 13(0.93) | 11(0.79) |
| S39 | 22.932940,113.113350,-5.53 | Lecong | Parklands | 2025.08.03 | 17 | 17(1) | 13(0.76) |
| S40 | 22.932940,113.113350,-5.53 | Lecong | Parklands | 2025.08.03 | 22 | 22(1) | 11(0.5) |
| S41 | 22.963381,113.143616,3.23 | Lecong | Parklands | 2025.08.04 | 19 | 19(1) | 18(0.95) |
| S42 | 22.968616,113.131584,4.26 | Lecong | Parklands | 2025.08.04 | 54 | 54(1) | 40(0.74) |
| S43 | 22.969074,113.119949,-0.59 | Lecong | Parklands | 2025.08.04 | 41 | 30(0.73) | 21(0.51) |
| S44 | 22.963629,113.131615,-7.78 | Lecong | Residences | 2025.08.04 | 8 | 8(1) | 5(0.63) |
| S45 | 22.965038,113.127007,-5.94 | Lecong | Parklands | 2025.08.04 | 47 | 47(1) | 28(0.6) |
| S46 | 22.959116,113.124023, -1.38 | Lecong | Residences | 2025.08.04 | 19 | 14(0.74) | 12(0.63) |
| S47 | 22.959806,113.119278,0.45 | Lecong | Residences | 2025.08.04 | 26 | 25(0.96) | 14(0.54) |
| S48 | 22.956518,113.128998,-1.48 | Lecong | Parklands | 2025.08.04 | 8 | 8(1) | 7(0.88) |
| S49 | 22.948227,113.131241,2.10 | Lecong | Parklands | 2025.08.04 | 14 | 9(0.64) | 9(0.64) |
| S50 | 22.938456,113.122124,-0.55 | Lecong | Parklands | 2025.08.04 | 23 | 18(0.78) | 11(0.48) |
| S51 | 22.952625,113.143990, -0.38 | Lecong | Parklands | 2025.08.05 | 28 | 28(1) | 18(0.64) |
| S52 | 22.949583,113.137161,-2.05 | Lecong | Residences | 2025.08.05 | 20 | 15(0.75) | 13(0.65) |
| S53 | 22.956423,113.132774,-1.59 | Lecong | Residences | 2025.08.05 | 37 | 36(0.97) | 21(0.57) |
| S54 | 22.934568,113.134209,-2.14 | Lecong | Parklands | 2025.08.05 | 1 | 1(1) | 1(1) |
| S55 | 22.933887,113.137299,-2.47 | Lecong | Parklands | 2025.08.05 | 11 | 9(0.82) | 8(0.73) |
| S56 | 22.931562,113.134583,-6.87 | Lecong | Parklands | 2025.08.05 | 12 | 12(1) | 10(0.83) |
| S57 | 22.929783,113.129494,-2.56 | Lecong | Parklands | 2025.08.05 | 8 | 8(1) | 7(0.88) |
| S58 | 22.927799,113.131149,3.75 | Lecong | Parklands | 2025.08.05 | 9 | 6(0.67) | 5(0.56) |
| S59 | 22.923870,113.141296,1.57 | Lecong | Parklands | 2025.08.05 | 5 | 4(0.8) | 3(0.6) |
| S60 | 22.927221,113.141434,0.54 | Lecong | Residences | 2025.08.05 | 12 | 12(1) | 3(0.25) |
| S61 | 22.958267,113.146805,3.02 | Beijiao | Parklands | 2024.08.06 | 35 | 33(0.94) | 13(0.37) |
| S62 | 22.952589,113.154701,-0.96 | Beijiao | Parklands | 2024.08.06 | 11 | 10(0.91) | 6(0.55) |
| S63 | 22.943665,113.147926,3.20 | Beijiao | Parklands | 2024.08.06 | 8 | 6(0.75) | 4(0.5) |
| S64 | 22.942019,113.145805,-7.25 | Beijiao | Parklands | 2024.08.06 | 31 | 30(0.97) | 9(0.29) |
| S65 | 22.934780,113.165100,-5.23 | Beijiao | Residences | 2024.08.06 | 16 | 16(1) | 6(0.38) |
| S66 | 22.931213,113.165192,4.31 | Beijiao | Residences | 2024.08.06 | 7 | 7(1) | 3(0.43) |
| S67 | 22.924730,113.158623,-1.30 | Beijiao | Parklands | 2024.08.06 | 37 | 34(0.92) | 14(0.38) |
| S68 | 22.917967,113.147591,2.80 | Beijiao | Parklands | 2024.08.06 | 47 | 46(0.98) | 24(0.51) |
| S69 | 22.921291,113.152603,1.91 | Beijiao | Residences | 2024.08.06 | 9 | 9(1) | 4(0.44) |
| S70 | 22.912149,113.161278,1.77 | Beijiao | Parklands | 2024.08.06 | 24 | 24(1) | 19(0.79) |
| S71 | 22.951502,113.176147,4.86 | Beijiao | Parklands | 2025.08.09 | 28 | 28(1) | 20(0.71) |
| S72 | 22.929945,113.196190,-5.67 | Beijiao | Parklands | 2025.08.09 | 141 | 138(0.98) | 82(0.58) |
| S73 | 22.919039,113.199074,-1.49 | Beijiao | Parklands | 2025.08.09 | 56 | 56(1) | 42(0.75) |
| S74 | 22.919886,113.204460,-6.68 | Beijiao | Parklands | 2025.08.09 | 18 | 17(0.94) | 16(0.89) |
| S75 | 22.903912,113.209663,-0.82 | Beijiao | Residences | 2025.08.09 | 79 | 78(0.99) | 57(0.72) |
| S76 | 22.901756,113.198616,-2.17 | Beijiao | Parklands | 2025.08.09 | 15 | 15(1) | 8(0.53) |
| S77 | 22.901625,113.180038,0.27 | Beijiao | Parklands | 2025.08.09 | 13 | 12(0.92) | 10(0.77) |
| S78 | 22.909956,113.180351,-7.29 | Beijiao | Residences | 2025.08.09 | 125 | 117(0.94) | 78(0.62) |
| S79 | 22.926691,113.174591,-17.54 | Beijiao | Parklands | 2025.08.09 | 28 | 25(0.89) | 23(0.82) |
| S80 | 22.938677,113.185883,-3.38 | Beijiao | Parklands | 2025.08.09 | 57 | 57(1) | 27(0.47) |
| S81 | 22.955221,113.184517, -2.29 | Beijiao | Parklands | 2025.08.10 | 30 | 28(0.93) | 20(0.67) |
| S82 | 22.947800,113.204201,-9.52 | Beijiao | Parklands | 2025.08.10 | 61 | 61(1) | 33(0.54) |
| S83 | 22.940449,113.226410,-9.86 | Beijiao | Parklands | 2025.08.10 | 72 | 69(0.96) | 44(0.61) |
| S84 | 22.934334,113.207855,-8.54 | Beijiao | Parklands | 2025.08.10 | 27 | 24(0.89) | 13(0.48) |
| S85 | 22.930031,113.224838,-11.53 | Beijiao | Parklands | 2025.08.10 | 6 | 3(0.5) | 1(0.17) |
| S86 | 22.937695,113.239998,5.73 | Beijiao | Parklands | 2025.08.10 | 50 | 47(0.94) | 23(0.46) |
| S87 | 22.969385,113.187569,-0.87 | Chencun | Parklands | 2025.08.10 | 88 | 81(0.92) | 60(0.68) |
| S88 | 22.982071,113.156708,-9.29 | Chencun | Parklands | 2025.08.10 | 38 | 37(0.97) | 20(0.53) |
| S89 | 22.972738,113.155708,-10.71 | Chencun | Parklands | 2025.08.10 | 4 | 3(0.75) | 3(0.75) |
| S90 | 22.975294,113.138359,-12.01 | Chencun | Parklands | 2025.08.10 | 47 | 46(0.98) | 25(0.53) |

